# *Pseudomonas aeruginosa* lectin LecB impairs keratinocyte fitness by abrogating growth factor signalling

**DOI:** 10.1101/629972

**Authors:** Alessia Landi, Muriel Mari, Tobias Wolf, Svenja Kleiser, Christine Gretzmeier, Isabel Wilhelm, Dimitra Kiritsi, Roland Thünauer, Roger Geiger, Alexander Nyström, Fulvio Reggiori, Julie Claudinon, Winfried Römer

## Abstract

Lectins are glycan-binding proteins with no catalytic activity and ubiquitously expressed in nature. Numerous bacteria employ lectins to efficiently bind to epithelia, thus facilitating tissue colonisation. Wounded skin is one of the preferred niches for *Pseudomonas aeruginosa*, which has developed diverse strategies to impair tissue repair processes and promote infection.

Here, we analyse the effect of the *P. aeruginosa* fucose-binding lectin LecB on human keratinocytes and demonstrate that it triggers events in the host, upon binding to fucosylated residues on cell membrane receptors, that extend beyond its role as an adhesion molecule. We found that LecB associates with several growth factor receptors and dampens their signalling pathways, leading to the arrest of cell cycle. Additionally, we describe a novel LecB-triggered mechanism to downregulate host cell receptors by showing that LecB leads to insulin-like growth factor receptor 1 internalisation, without receptor activation, and subsequent missorting towards intracellular endosomal compartments.

Overall, these data highlight that LecB is a multitask virulence factor that, through subversion of several host pathways, has a profound impact on keratinocyte proliferation and survival.

## Introduction

Bacteria can employ many different strategies to infect host cells. In all cases the initiation of an infection requires the recognition of specific structures on the host cell plasma membrane. This is often achieved by lectins, which bind to glycosylated residues on proteins or lipids present on the cell surface, mediating the attachment of the bacterium to the cell. Multivalency is an important feature of most lectins. On one hand, multivalency increases the binding affinity and specificity of the lectin-glycan interaction[1]. On the other hand, the binding of lectins to multiple cell surface receptors can induce receptor clustering and plasma membrane rearrangement, triggering their entry into the host[2–4].

*Pseudomonas aeruginosa* is a Gram-negative bacterium, ubiquitously spread in nature. It is an opportunistic pathogen that can cause severe infections, especially in immunocompromised individuals, due to its resistance to most of the available antibiotics and its ability to form impenetrable biofilms. Hence, it has been classified in the “priority 1/critical” category of the World Health Organisation (WHO) global priority pathogens list (global PPL) of antibiotic-resistant bacteria to promote the research and development of new antibiotic treatments[5].

It is frequently implicated in hospital-acquired infections, where it has been reported to cause different types of infections. Wounded skin, following traumatic injuries, surgery or burns, is one of the preferentially targeted tissue by the bacterium, which has also been associated with the delay and prevention of wound healing. The presence of *P. aeruginosa* correlates in fact with a bad prognosis of healing and leads to the persistence of the inflammatory stage of the wound healing process[6,7].

*P. aeruginosa* possesses two tetravalent lectins in its arsenal of virulence factors, LecA and LecB (also called PA-IL and PA-IIL, respectively). LecB is a tetramer, consisting of four monomers with high specificity for L-fucose and its derivatives[8,9]. LecB production is regulated by rhl and *Pseudomonas* quinolone signal (PQS), part of the quorum sensing systems[10,11]. Once synthesised, it is exposed on the outer bacterial membrane, where it has been described to interact with the outer membrane porin OprF[12,13].

The current assumption is that LecB mainly functions by promoting the adhesion of the bacterium to both the host cell and the exopolysaccharide matrix, which encases bacterial cells together. However, several *in vitro* and *in vivo* studies have shown LecB to act not only as an adhesin, but also as an important virulence factor, capable of triggering additional host cell responses[14,15]. LecB has been reported to be a determinant of *P. aeruginosa* cytotoxicity in lung epithelial cells and to block ciliary beating in human airways[16,17]. LecB-negative mutant bacteria exhibit an impaired biofilm formation in comparison with wild-type strains and no type VI pili assembly[12,18]. Furthermore, LecB induced alveolar capillary barrier injury *in vivo*, leading to a higher bacterial dissemination into the bloodstream[17]. Previous studies have described additional effects of LecB in inhibiting cell migration and proliferation[19]. However, its precise mechanism of action has not yet been elucidated and none of the existing studies have addressed its role in skin infections.

Here, we report that the *P. aeruginosa* lectin LecB is present in human chronically infected wounds, implying its contribution to the persistence of wound infections. Moreover, we show that insulin like growth factor-1 receptor (IGF-1R) co-precipitates with LecB and that LecB leads to IGF-1R internalisation and missorting towards intracellular LC3-positive compartments. Notably IGF-1R is internalised without being activated. We further demonstrate that LecB blocks the cell cycle and induces cell death preceded by a strong vacuolisation. These vacuoles, which possess peculiar morphological features, originate from ruffle-like structures at subdomains of the plasma membrane where LecB is enriched. Therefore, we propose that LecB, in addition to play a role as an adhesion factor, both misregulates growth factor receptor signalling and subverts the endocytic system, leading to the impairment of vital keratinocyte functions.

## Results

### LecB is present in chronically infected human wounds

Although LecB is enriched in biofilms, which often characterise chronic wounds, no evidence had yet been shown of its presence in wounds. Before we addressed the effects of LecB on human keratinocytes on a molecular level, we first verified the presence of LecB in infected human wounds. To this aim, we collected chronic wounded tissue from patients infected with *Pseudomonas aeruginosa*, as shown by wound swabs. We stained the paraffin-embedded tissue sections with an antibody against *P. aeruginosa* to confirm its presence in the wounds. As control, we used normal skin samples. Indeed, we could detect the presence of *P. aeruginosa*, either as biofilm or in the form of small colonies (Fig. 1A). We subsequently stained for LecB and strikingly, we found this lectin to be distributed in the wound sections, both in the keratinocyte layers and in the dermis (Fig. 1B). In contrast, no LecB was found in the normal skin control sample, proving the specificity of the antibody. This result provides the first evidence of LecB in chronic human wounds, with the superinfection possibly playing a role in the wound chronicity.

**Fig. 1:**
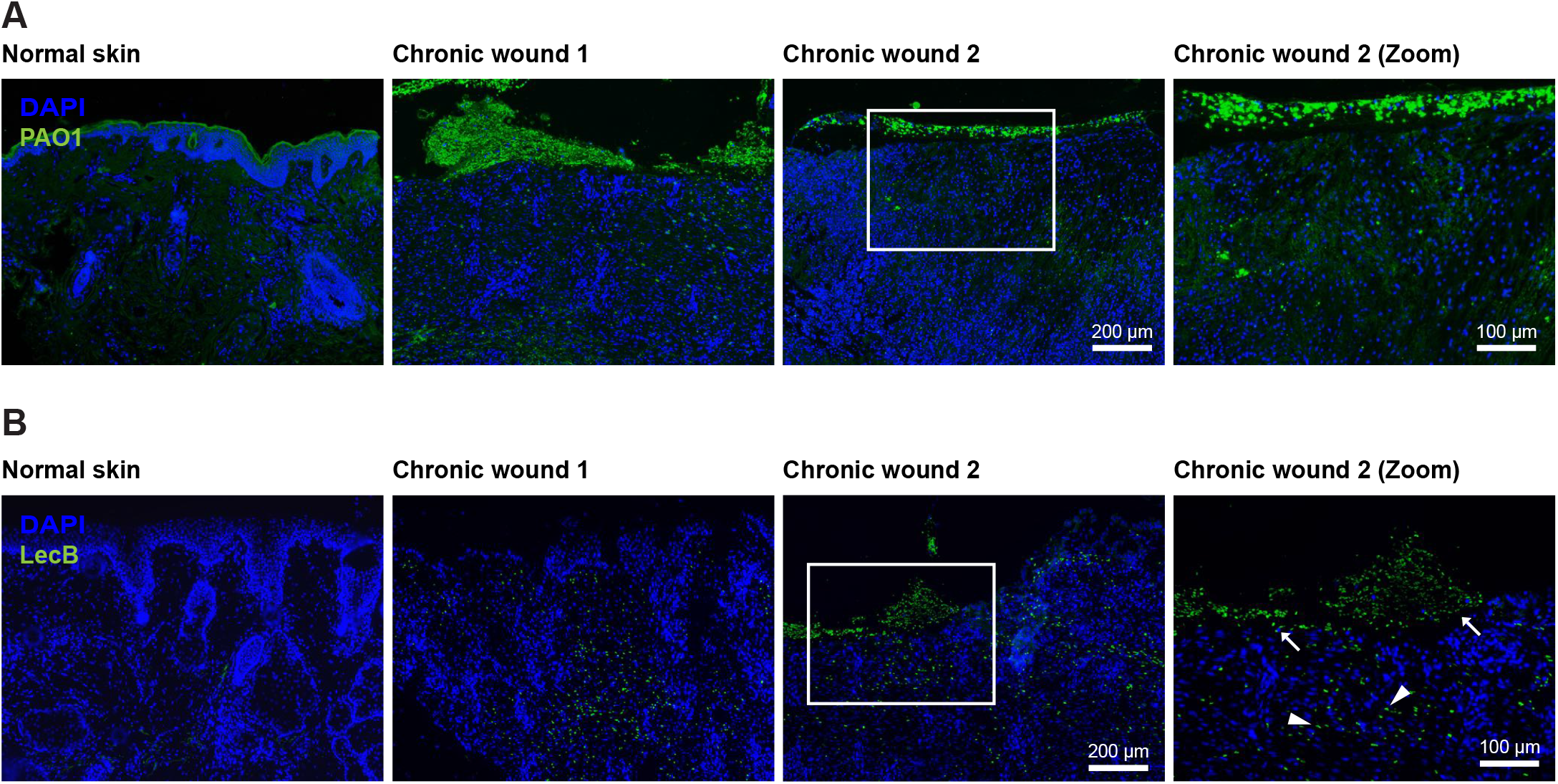
LecB localisation in chronically infected human wounds. Tissue sections of human infected wounds embedded in parafffin and stained for (A) Pseudomonas aeruginosa (green) and for (B) LecB (green). Normal skin is used as negative control. Rectangular squares refer to the zoomed area. Arrows point at LecB localised in the epidermal layers; arrowheads indicate LecB distributed in the dermis.

### LecB co-precipitates with essential plasma membrane receptors in normal human keratinocytes

Next, in order to study the mechanism of action of LecB in molecular detail, we moved to normal human keratinocytes (NHKs), the predominant cell type in the epidermis. LecB plays a crucial role in the adhesion of the bacterium to host cells[17], implying the necessity for the lectin to target plasma membrane receptors via their glycosylated residues. Therefore, we screened for potential LecB interacting proteins via a pull-down assay. Briefly, we cultured NHKs in presence or absence of 5 μg/ml (106 nM) of biotinylated LecB, lysed them under conditions that preserve protein-protein interactions and incubated with streptavidin agarose beads to precipitate LecB in complex with interacting proteins. We performed on bead digestion and analysed the obtained peptides by protein mass spectrometry. This analysis revealed the presence of important cell growth factor receptors within the enriched proteins in the LecB pull down fractions (SI Appendix, Table S1). The mass spectrometry results were verified by western blot, confirming the presence of two of the major keratinocyte growth factor receptors implicated in epidermal keratinocytes proliferation and migration, the insulin-like growth factor receptor 1 (IGF-1R) and the epidermal growth factor receptor 1 (EGF-1R)[20,21], in the eluted fractions. In addition to stimulating the cells with biotinylated LecB, an excess of L-fucose (30 mM) was added to clarify whether the interaction with IGF-1R or EGFR-1R was carbohydrate-dependent. As L-fucose saturates the carbohydrate-binding pockets of LecB, no binding to cell membranes could be detected when LecB and L-fucose were simultaneously applied to the cells as neither IGF-1R or EGF-1R, nor LecB could be precipitated (Fig. 2A and SI Appendix, Fig. S1A).

**Fig. 2:**
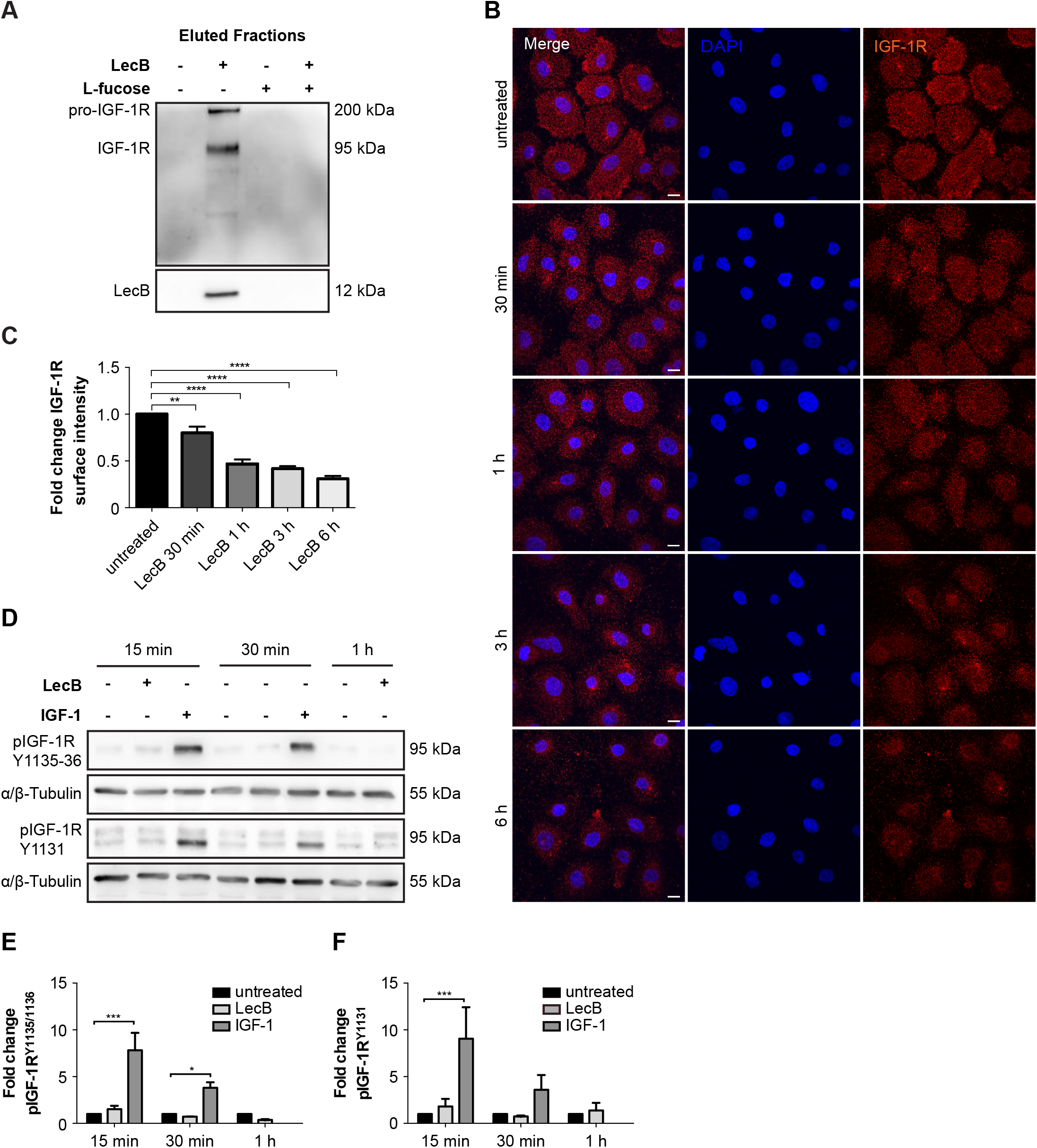
LecB depletes IGF-1R from the plasma membrane without inducing its activation. (A) Western Blot of eluted samples from pull-down assay. Normal human keratinocytes (NHKs) were stimulated in presence of 5 μg/mL biotinylated LecB (106 nM), medium, L-fucose (30 mM) and biotinylated LecB + L-fucose. The lysates were incubated with streptavidin beads and further eluted. Western Blot was performed and IGF-1R and LecB were detected in the precipitated fractions. (B, C) Surface staining of keratinocytes treated with LecB (5 μg/mL) for the indicated time points. (B) Images show maximum intensity projections. After stimulation cells were stained for IGF-1R (red) and DAPI (blue). Scale bar: 10 μm. (C) Quantification of IGF-1R surface intensity from n = 4 independent experiments. Bars display the mean value ± standard error mean. (D) Representative blots from lysates after LecB (5 μg/mL) and IGF-1 (100 ng/mL) stimulation for the indicated times. Antibody against different phosphorylation sites of IGF-1R (Tyr1131 and Tyr1135/1136) were used. Tubulin was employed as loading control. (E, F) Blot quantification. Phosphorylated IGF-1R levels are represented as fold change compared to the loading control from n = 3 independent experiments. *denotes p < 0.05; ** denotes p < 0.01; *** denotes p < 0.001; **** denotes p < 0.0001; ns denotes not significant.

We further investigated in detail the effect of LecB on insulin-like growth factor receptor 1 (IGF-1R), due to its higher fold enrichment. We hypothesised that LecB interaction with IGF-1R, may trigger receptor internalisation. Indeed, surface staining experiments showed that IGF-1R is depleted from the plasma membrane of keratinocytes in a time-dependent manner as a consequence of LecB incubation (Fig. 2B, C). IGF-1R internalisation from the cell surface was also visible after stimulation with 100 ng/ml of insulin-like growth factor 1 (IGF-1) (SI Appendix, Fig. S2A, B). However, while IGF-1 induced a fast autophosphorylation of the receptor on multiple tyrosine residues (Tyr1135, Tyr1131 and Tyr1136), LecB did not (Fig. 2D, E, F). Thus, we concluded that LecB does not lead to the activation of IGF-1R, but rather triggers a “silent” receptor internalisation.

### LecB impairs cell survival signalling pathways and leads to cell cycle arrest

Growth factor receptors activate two major downstream signalling pathways, the MAPK/ERK and the PI3K-AKT pathways, which are responsible for inhibition of apoptosis, stimulation of protein synthesis and cell proliferation[22,23]. While the former ultimately results in the phosphorylation and activation of extracellular signal-regulated kinase 1/2 (ERK1/2), the latter activates the mammalian target of rapamycin (mTOR). In order to check whether these two signalling cascades were affected by LecB, we monitored ERK1/2 and mTOR phosphorylation at Thr202/Tyr204 and Ser2448, respectively. The amounts of phosphorylated proteins were normalised to the respective pan-proteins. Indeed, LecB did not lead to the phosphorylation of both ERK1/2 and mTOR. ERK1/2 phosphorylation actually significantly decreased 1 h post stimulation (Fig. 3A-D). In contrast, LecB activated 5’ adenosine monophosphate-activated protein kinase (AMPK), a cellular energy and nutrient status sensor, activated in response to cellular energy depletion (Fig. 3A, E). AMPK phosphorylation significantly increased after 3 h post stimulation and was inhibited by addition of L-fucose, confirming to be LecB-dependent (SI Appendix, Fig. S3A, B).

**Fig. 3:**
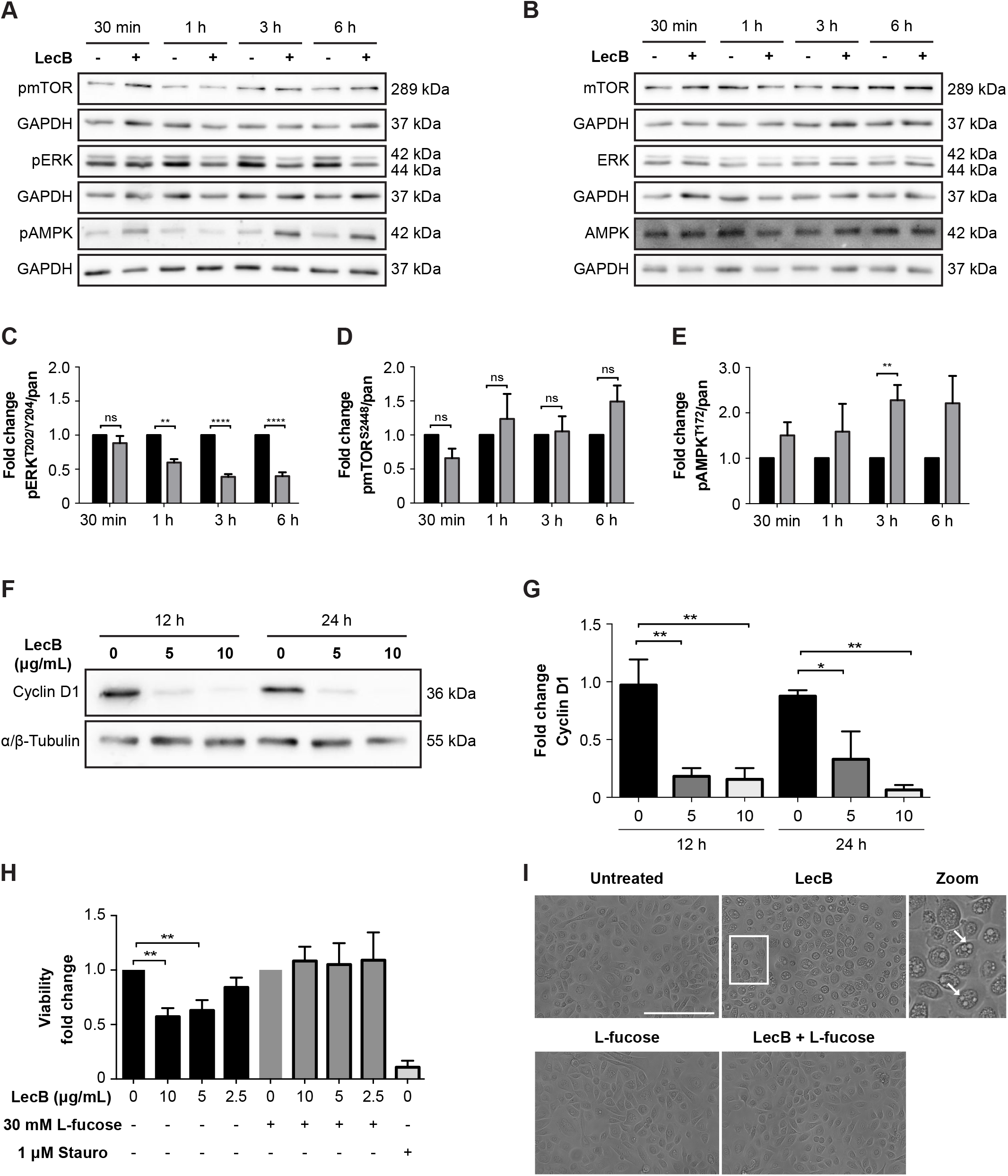
LecB impairs cell survival signalling pathways and leads to cell cycle arrest. (A-E) Representative blots and relative quantifications of N = 3 independent experiments. NHKs were treated with 5 μg/mL LecB for the indicated time and lysates were subjected to SDS-PAGE and Western Blot analysis using the designated anti-phospho (A) and anti-pan antibodies (B). GAPDH was added as loading control. Graphs (C-E) depict the fold change of the phosphorylated protein compared to pan levels and represent the mean value ± standard error mean. (F) Western Blot showing cyclin D1 levels after 12 and 24 h of LecB stimulation (5 or 10 μg/mL). (G) Quantification of cyclin D1 relative to tubulin. The mean value ± standard error mean of N = 3 independent experiments is reported. (H) MTT assay assessing the cytotoxic effect of LecB. NHKs were treated with the indicated LecB concentration with or without 30 mM L-fucose for 24 h. Staurosporin (1 μM) was employed as positive control. MTT was added to the medium and left for 4 h. The absorbance at 570 nm was measured and plotted as fold change compared to the untreated or L-fucose treated sample. The mean value ± standard error mean of N = 5 is plotted. (I) Representative images of keratinocyte monolayers after 24 h exposure to 5 μg/mL LecB with or without 30 mM L-fucose. White arrows in the zoomed image point at vacuolar structures induced by lectin treatment. Scale bar: 200 μm. * denotes p < 0.05; ** denotes p < 0.01; *** denotes p < 0.001; **** denotes p < ns denotes not significant.

The activation of AMPK correlates with the inhibition of ERK1/2, since the latter has been reported to negatively regulate the phosphorylation of AMPK via the liver kinase B1 (LKB1)[24,25]. Furthermore, consistently with the inhibition of ERK1/2 activity, we observed a strong cyclin D1 degradation (Fig. 3F, G), which led to the arrest of cell cycle and thus to the reduction of cell viability (Fig. 3H). Interestingly, the cytotoxic effect was preceded by an extensive cytoplasmic vacuolisation (Fig. 3I). L-fucose supplementation rescued the cells and restored viability (Fig. 3H, I). Altogether, these observations elucidate that LecB impairs a key mechanism that regulates cell survival and proliferation, resulting in the blockage of cell cycle.

### The cytotoxic effect of LecB is accompanied by the formation of intraluminal vesicles-containing vacuoles where LecB is localised

Next, we used transmission electron microscopy (TEM) to further investigate the cellular vacuolisation triggered by LecB treatment. Untreated keratinocytes present several light content vacuoles (Category 1), corresponding to endo-lysosomal compartments (Fig. 4A), LecB treated cells display numerous degradative vesicles (Category 2) and an additional type of vacuoles (Category 3), with irregular shapes and variable sizes (Fig. 4B). Surprisingly, these vacuoles contain a large number of clearly defined intraluminal vesicles (Fig. 4B) and they seem to cluster together to form an intricate network (Fig. 4B’, zoom). To get insights into the origin of these structures, we performed a time-course experiment by collecting cells at different time points, from 30 min to12 h after exposure to LecB (SI Appendix, Fig. S4A-E). The Category 3 vacuoles were detectable after 30 min of LecB incubation, but they were quite rare. However, they became more prominent 3 h post LecB exposure (SI Appendix, Fig. S4D). Interestingly, specific regions of the plasma membrane of the LecB-treated keratinocytes were extremely irregular and characterised by the presence of ruffle-like structures (SI Appendix, Fig. S4A-E). As the number of Category 3 vesicles located at these regions increased over time, we hypothesised that the intraluminal vesicles-containing vacuoles originate from subdomains of the plasma membrane upon LecB interaction with surface proteins. This notion implies that LecB is present in the Category 3 vacuoles. To test this hypothesis, we exposed keratinocytes to biotinylated LecB before performing immuno-electron microscopy (IEM) using anti-biotin antibodies (Fig. 4C-F). Notably, we specifically detected LecB on both the limiting membrane of the Category 3 vacuoles and to a minor extent on their intraluminal vesicles (Fig. 4D). Moreover, we also identified biotinylated LecB at the plasma membrane, where it was enriched in the ruffle-like regions (Fig. 4F), further indicating that LecB drives the formation of a plethora of vesicles originating from the cell plasma membrane.

**Fig. 4:**
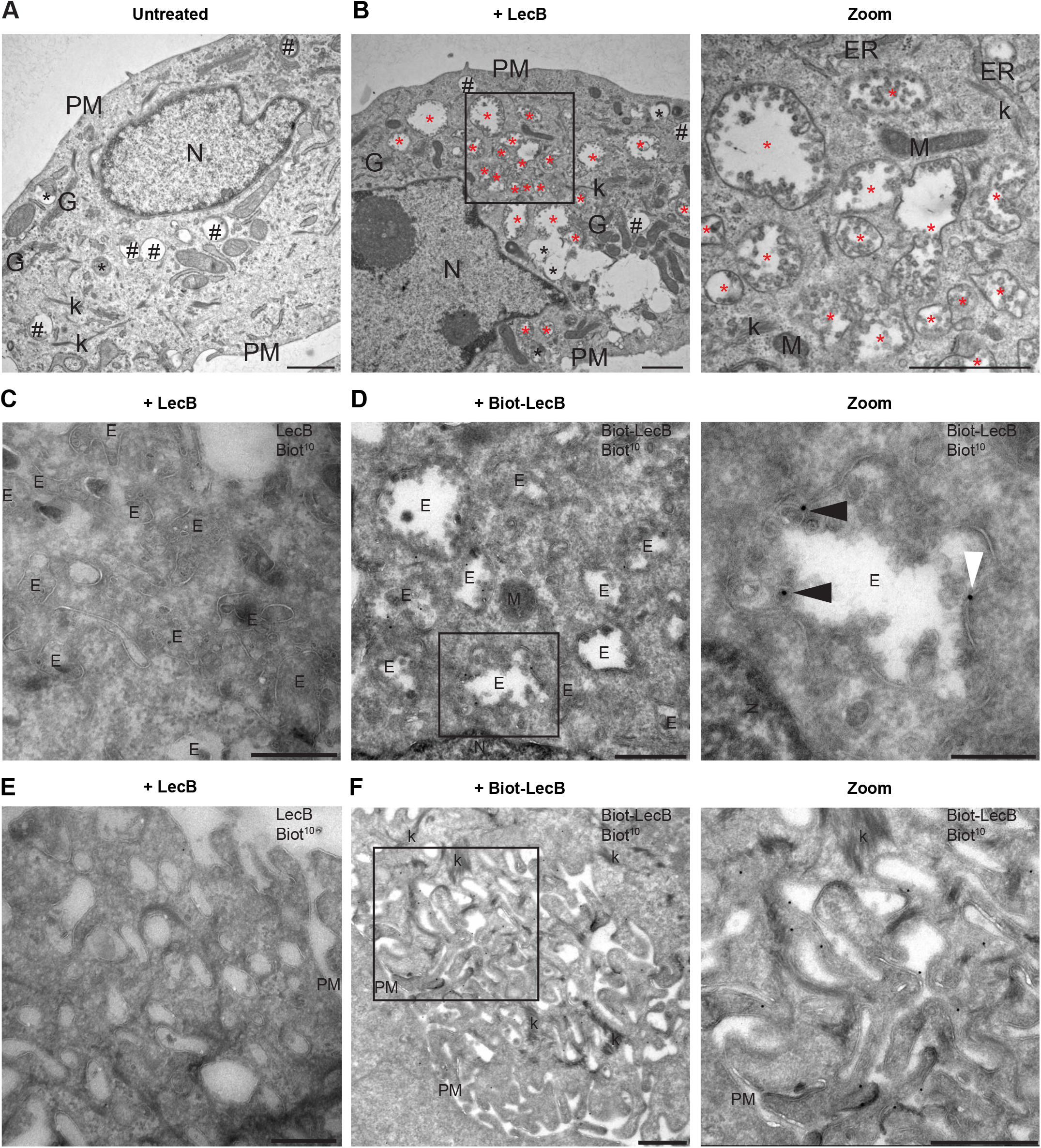
The cytotoxic effect of LecB is preceded by the formation of intraluminal vesicles containing vacuoles. (A-B) NHKs were incubated with 5 μg/mL LecB and processed for conventional electron microscopy at 12 h post incubation. Representative electrographs of the untreated (A) and LecB treated (B) cells. Black hashtags point at category 1 vacuoles, while black and red asterisks indicate category 2 and category 3 vacuoles, respectively. Zoomed image of panel B shows a higher magnification of the category 3 vacuoles containing intraluminal vesicles. Scale bars: 1 μm. (C-F) Cells treated with 5 μg/mL biotinylated LecB or un-tagged LecB for 12 h were subjected to immuno-EM. (C, E) Control staining showing no aspecific antibody binding. (D) LecB localises in internal vesicles, either in the lumen (panel D zoom, black arrowheads) or at the limiting membrane (white arrowheads). (F) LecB localisation at the plasma membrane, in the ruffle-like region. Scale bars: 500 nm. Scale bar zoom panel D: 200 nm. E: endosome; ER: endoplasmic reticulum; k: keratin; G: Golgi apparatus; M: mitochondrion; N: nucleus; PM: plasma membrane.

### LecB is trafficked towards an endocytic degradative route

Next, we proceeded to characterise the intracellular LecB-positive vacuoles detected by IEM in immunofluorescence experiments, where keratinocytes exposed to fluorescently-labelled LecB were stained with antibodies recognising different organelle marker proteins. We found a time-dependent colocalisation of LecB with microtubule-associated proteins 1A/B light chain 3B (LC3) and ras-related protein Rab-9 (RAB9A), which are generally enriched on autophagosomes and late endosomes, respectively (Fig. 5A, B). Moreover, but to a minor extent, we also observed a time-dependent colocalisation between LecB and lysosomal-associated membrane protein 1 (LAMP1) (SI Appendix, Fig. S5A). In contrast, no colocalisation with the recycling endosome marker protein RAB11 was detected (SI Appendix, Fig. S5B). Endocytosis and internalisation of LecB was also assessed biochemically by analysing the ubiquitination of the proteins associated with LecB, as a large number of growth factor receptors undergo ubiquitination upon endocytosis[26,27]. In a time-course pull-down experiment we detected a decrease of the total levels of IGF-1R and an increase of the overall cell ubiquitin levels in the eluted fraction, indicating that on one hand IGF-1R is targeted for degradation and on the other hand that LecB interacting partners are ubiquitinated, thus very likely internalised (SI Appendix, Fig. S6A, B).

**Fig. 5:**
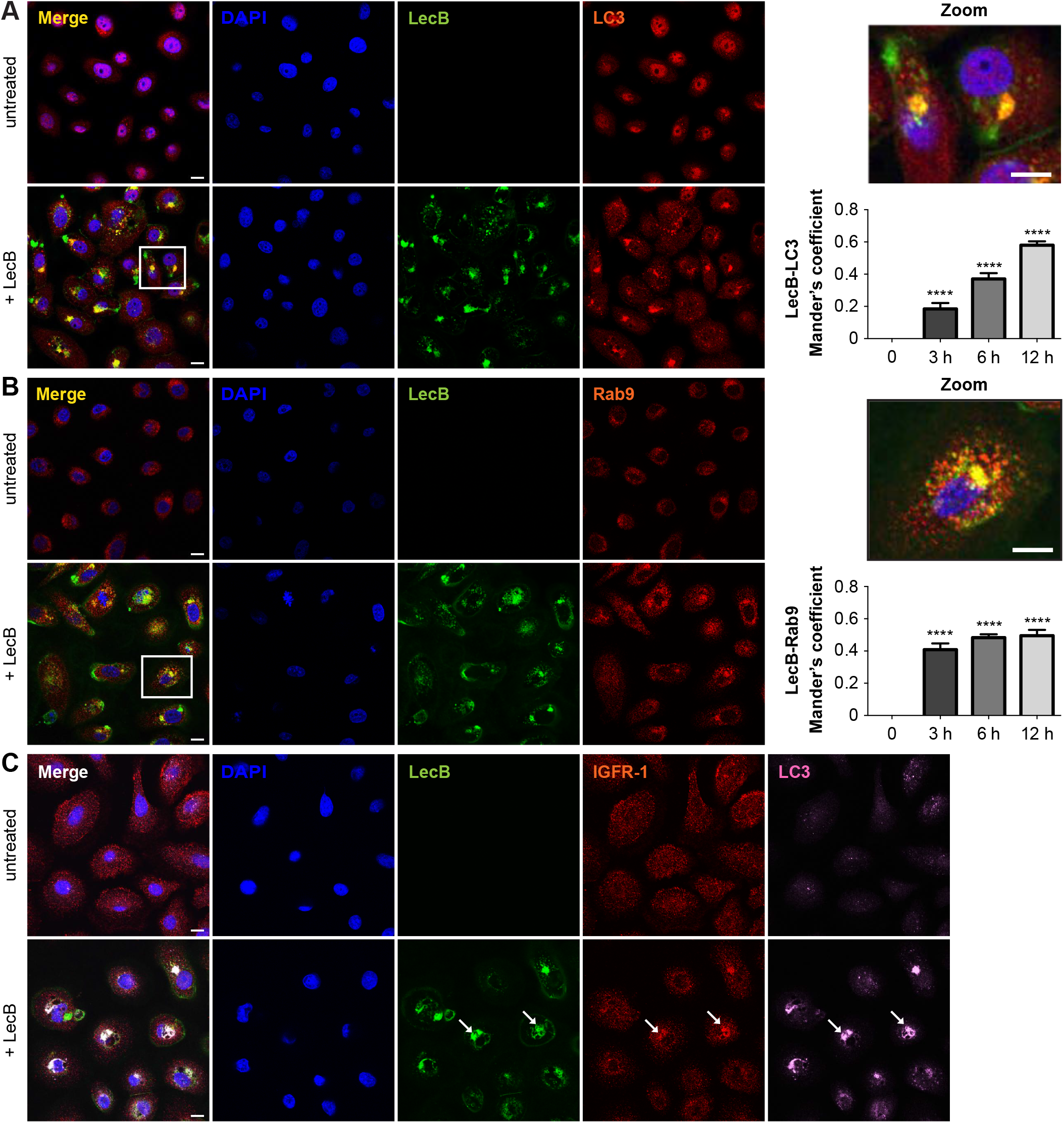
LecB colocalises with LC3, Rab9 and IGF-1R. (A, B) Confocal micrographs with respective colocalisation quantification of keratinocytes stimulated with 5 μg/mL of fluorescently labelled LecB (green). Panels indicated as “+ LecB” refer to 12 h incubation. After fixation and permeabilisation, the cells were stained for LC3 (A) and Rab9 (B) (both in red). Graphs report the mean value ± standard error mean of Mander’s overlap coefficients calculated from at least 3 independent experiments. Scale bar: 10 μm. **** denotes p < 0.0001. (C) Confocal micrographs of keratinocytes treated with 5 μg/mL of fluorescently labelled LecB (green) and stained for LC3 (pink) and IGF-1R (red). Panel indicated as “+ LecB” refers to 12 h incubation. White arrows point at colocalization among LecB, LC3 and IGF-1R. Scale bar: 10 μm. N = 3.

As we saw a time-dependent increase in colocalisation between LC3 and LecB structures (Fig. 5A), we decided to investigate this further. LC3 is an ubiquitin-like protein that upon autophagy induction is converted from a cytoplasmic form (i.e., LC3-I) to one associated to the autophagosomal membranes (i.e, LC3-II) through conjugation to phosphatidylethanolamine[28]. LC3-II can also be generated on the limiting membrane of endosomes during processes like LC3-associated phagocytosis (LAP)[29,30]. The fact that LecB was detected on single membrane vacuoles (autophagosomes are double-membrane vesicles with cytoplasmic content) that are positive for RAB9 (Fig. 4 and 5B), inferred in an activation of LAP. Although induction of both autophagy and LAP induces the formation of LC3-II, only autophagy leads to the degradation of this conjugate[31]. Indeed, we observed a time-dependent increase of LC3-II levels upon exposure of keratinocytes to LecB (SI Appendix, Fig. S7A, B). However, we did not detect a turnover of this protein since no difference in LC3-II levels could be observed when LecB-treated keratinocytes were incubated with either bafilomycin A1, a lysosomal inhibitor, or cycloheximide to assess LC3-II stability (SI Appendix, Fig. S7C-F).

Finally, we investigated whether IGF-1R also colocalised with LecB-/ LC3-positive vesicles. Notably, while we found colocalisation between IGF-1R, LecB and LC3, we did not observe LC3 recruitment when cells were treated exclusively with IGF-1 (Fig. 5C, SI Appendix, Fig. S5C). Taken together these results demonstrate that LecB is trafficked towards degradative compartments, leading to missorting of IGF-1R. They also suggest that LAP participates in targeting LecB-IGF-1R complexes to degradation.

## Discussion

Bacterial lectins have predominantly been described as adhesion proteins, which promote the attachment of the bacterium to cell plasma membrane through their interaction with host cell surface glycans[32]. This study provides new insights into the role of *P. aeruginosa* lectin LecB as virulence factor that impairs growth factor receptor signalling and trafficking, thereby affecting keratinocyte fitness. Pull-down studies showed the co-isolation of LecB with several plasma membrane receptors in keratinocytes, among which IGF-1R and EGF-1R were two of the principal interactors. We further demonstrated that LecB depletes IGF-1R from the plasma membrane by inducing a time-dependent receptor internalisation. To modulate signalling, IGF-1R (as well as EGF-1R) needs to be activated by ligand binding and consequent autophosphorylation on three tyrosine residues (i.e, Tyr1135, Tyr1131 and Tyr1136) in the C-terminal tail[33,34]. Interestingly, unlike IGF-1, LecB mediates receptor internalisation without receptor activation and triggers IGF-1R endocytic trafficking towards degradative compartments.

PI3K/AKT and MAPK/ERK signalling cascades have been intensively investigated for their role in promoting cell and organismal growth upon growth factor signals. This is mainly achieved by AKT-mediated activation of mTOR, which is a key regulator of anabolic processes necessary during cell growth[35,36] and by the phosphorylation of ERK1/2, which results in cell proliferation[37,38]. Indeed, we found that neither mTOR nor ERK1/2 was activated upon LecB. Specifically, we observed a decrease in the levels of phospho-ERK1/2 after 1 h of LecB exposure, which correlates with the increase of phospho-AMPK. Reciprocal feedback loops have been described to operate between the AMPK and the MAPK/ERK pathways and, whereas ERK1/2 activation was reported to be attenuated upon stimulation of WT mouse embryonic fibroblasts with the AMPK activator A-769662, little effect on ERK1/2 phosphorylation was detected when AMPK-null MEFs were used[39,40]. The suppression of cell survival signalling pathways can also clarify the arrest of the cell cycle in the G1/S transition observed upon LecB incubation. ERK1/2 activity is in fact required for both the cell cycle entry and the suppression of negative cell cycle regulators. Specifically, cyclin D1 expression is enhanced by the activation of the MAPK/ERK pathway[41,42], via the transcription factors c-Fos[43] and Myc[44]. Cells entering into the S phase show low levels of cyclin D1, which is again induced and reaches high levels to ensure the progression into the G2 phase[45]. LecB stimulation mediates a strong reduction of cyclin D1, whose levels are not restored over time, resulting in the arrest of the cell cycle and subsequently in the induction of cell death. Therefore, we speculate that LecB, despite devoid of catalytic activity, may contribute in promoting tissue damage to facilitate bacterial dissemination and extracellular multiplication during wound infections. Furthermore, it is possible that LecB induces a similar fate to other cell types, including immune cells, thus promoting the infection. Nonetheless, future work will provide new insights on the impact of LecB on the immune response during chronic wound infections.

Interestingly, the cytotoxic effect of LecB was preceded by an extensive cytoplasmic vacuolation. Electron microscopy inspection of LecB-treated cells revealed the peculiar nature of these vacuoles that over time accumulate and display an increased number of intraluminal vesicles. We also show that these vacuoles originate from ruffle-like plasma membrane regions where LecB was also found to be enriched. In the case of other bacterial lectins, such as the B-subunit of Shiga toxin (StxB) from *Shigella dysenteriae* and LecA from *Pseudomonas aeruginosa*, the sole interaction with the glycosphingolipid receptor globotriaosyl ceramide (Gb3) is sufficient to drive membrane shape changes leading to their subsequent uptake[3,46]. In the case of LecB, however, pull-down experiments point to the existence of several receptors, also attributable to the fact that fucosylation is a very common modification. Additionally, LecB exists as a tetramer and each monomer possesses a binding pocket with the highest affinity to L-fucose and its derivatives[8,47]. Although the L-fucose dissociation constant is in the micromolar range (2.9 μM), an increase in avidity can be achieved by a higher degree of interactions. This implies that LecB might crosslink several different surface receptors, thus inducing a higher degree of membrane rearrangements that could explain the extensive alterations at the plasma membrane. Moreover, multiple interactions provide the bacterium with additional resources for the initiation of host tissue colonisation.

By following LecB trafficking we sought to characterise the nature of LecB-mediated vacuoles, which, from a structural point of view, shows similarities with multivesicular bodies (MVBs), as they contain numerous intraluminal vesicles. Immunofluorescence experiments revealed a time-dependent increase of colocalisation between LecB and Rab9, LC3 and LAMP1. Therefore, our data indicate that LC3 is recruited on LecB containing late endosomes, which may further favour their lysosomal degradation through a process that may be similar or identical to LAP. LAP can be triggered by several receptors. In addition to toll-like receptors (TLRs), whose signalling during phagocytosis rapidly recruits LC3 to phagosomes, the phosphatidylserine receptor TIM4 or the C-type lectin Dectin-1 can also induce LAP to efficiently clear dead cells and to facilitate antigen presentation, respectively[29,48,49]. LC3 can associate with single-membrane phagosomes even in absence of pathogens or dead cells, such in the case of phagocytosis and degradation of photoreceptor outer segments (POS) by retinal pigment epithelium or during the secretion of mucins in goblet cells[50,51]. Our data indicate that IGF-1R colocalises with LecB and LC3, suggesting that LAP is involved in IGF-1R sorting for degradation, following LecB-mediated receptor internalisation.

The current model describes that tyrosine kinase receptors, upon ligand binding, are internalised and trafficked to early endosomes. From here, they can either be sorted to lysosomal degradation via MVBs or they can be recycled back to the plasma membrane. The equilibrium between IGF-1R degradation and recycling is essential to modulate receptor signalling[52,53]. Our data demonstrate that IGF-1R trafficking is subverted upon LecB, which preferentially sorts the receptor towards degradative routes, without activating it. We do not know yet whether there is a direct or indirect interaction between LecB and IGF-1R, despite being co-precipitated as complex. However, since IGF-1R is not the sole protein interacting with LecB in keratinocytes, we speculate that this “targeting by degradation” strategy can be valid for other membrane receptors as well (e.g., EGF-1R) and that it can be employed by *P. aeruginosa* to silence host cell receptors to favour tissue colonisation in wounds.

Wound infections represent a socio-economic burden for the healthcare system and, given that *P. aeruginosa* is very hard to eliminate with the available antibiotics, there is urgent need for the development of alternative therapeutic strategies[54,55]. Taken together, our findings shed new light on the *P. aeruginosa* lectin LecB, showing that it is capable of inducing a succession of cellular events, despite being devoid of catalytic activity and by virtue of its sole capability to bind to sugar moieties on the plasma membrane, and can set the basis for a better understanding of *P. aeruginosa* wound infections.

## Materials and Methods

### Antibodies, inhibitors and activators

Used antibodies are listed in table S2 and S3. Aprotinin (0.8 μM), leupeptin (11 μM) and pefabloc (200 μM), used as protease inhibitors, and phosphatase inhibitor cocktail 3 (1:100) were from Sigma Aldrich. To block LecB binding to the host cell membranes, L-fucose (Sigma Aldrich) was used at a concentration of 30 mM. Cycloheximide (20 μg/mL) and staurosporin (500 nM) were both purchased from Sigma Aldrich and used as protein synthesis inhibitor and as positive control for cell death, respectively. Bafilomycin was from InvivoGen and used at a concentration of 100 nM or 200 nM. Human insulin-like growth factor 1 (IGF-1) was from Thermo Fisher and used as IGF-1 receptor activator.

### Cell culture

Normal human keratinocytes (NHKs), kindly provided by D. Kiritsi (Department of Dermatology, Freiburg, Germany), were grown at 37°C and in the presence of 5 % CO_2_ in keratinocyte medium supplemented with bovine pituitary extract, epidermal growth factor and 0.5 % penicillin/streptomycin. Cells were seeded 24 h prior to each experiment. If not stated differently, all treatments were performed in complete medium.

### LecB production and *in vitro* labelling

Recombinant LecB (UniProt. ID: Q9HYN5_PSEAE) was produced from *E. coli* BL21(DE3) containing pET25pa21 and purified via chromatography using a mannose-agarose column as previously described (Mitchell et al. 2002; Chemani et al. 2009). The elute was dialysed in water for 7 days and finally lyophilised. The obtained powder was dissolved in phosphate-buffered saline (PBS, 138 mM NaCl, 2.7 mM KCl, 8 mM Na_2_HPO_4_, 1.5 mM KH_2_PO_4_, pH 7.4) and sterile filtered. Additionally, to exclude endotoxin contamination, LecB was further purified using an endotoxin removal column (Hyglos) and a LAL chromogenic endotoxin quantitation kit (Thermo Scientific) was carried out. The purity of the lectin was confirmed by SDS-PAGE. If not stated differently, LecB was used at a final concentration of 5 μg/ml (106 nM).

LecB was fluorescently labelled using Cy3 (GE Healthcare) or Alexa Fluor488 (Thermo Scientific) mono-reactive NHS ester and purified using Zeba Spin desalting columns (Thermo Scientific) according to the manufacturers’ instructions.

Biotinylated LecB was obtained using sulfo-NHS-SS-Biotin (Thermo Scientific) according to the supplier’s instructions and dialysed two times for 1 h in water and one time overnight in PBS.

### Mass spectrometry analyses

Cells were seeded in T-75 flasks and stimulated with biotinylated LecB at a concentration of 5 μg/mL. Upon treatment, cells were washed in phosphate-buffered saline (PBS) and lysed with 50 mM Tris-HCL pH 7.5, 150 mM sodium chloride, 1 % (v/v) IGEPAL CA-630 and 0.5 % (w/v) sodium deoxycholate in water. The protein lysates were incubated over night at 4 °C with streptavidin agarose beads (Thermo Scientific) to precipitate LecB-biotin-protein complexes. The beads were washed 4 times in PBS and then re-suspended in 8M urea. Immunoprecipitates were pre-digested with LysC (1:50, w/w) followed by reduction of disulfide bonds with 10 mM DTT and subsequent alkylation with 50 mM iodacetamide. Then samples were diluted 1:5 with ammonium bicarbonate buffer pH 8 and trypsin (1:50, w/w) was added for overnight digestion at RT. The resulting peptide mixtures were acidified and loaded on C18 StageTips[56]. Peptides were eluted with 80% acetonitrile (ACN), dried using a SpeedVac centrifuge, and resuspended in 2% ACN, 0.1% trifluoroacetic acid (TFA), and 0.5% acetic acid.

Peptides were separated on an EASY-nLC 1200 HPLC system (Thermo Fisher Scientific, Odense) coupled online to a Q Exactive HF mass spectrometer via a nanoelectrospray source (Thermo Fisher Scientific). Peptides were loaded in buffer A (0.5% formic acid) on in house packed columns (75 μm inner diameter, 50 cm length, and 1.9 μm C18 particles from Dr. Maisch GmbH). Peptides were eluted with a non-linear 270 min gradient of 5%–60% buffer B (80% ACN, 0.5% formic acid) at a flow rate of 250 nl/min and a column temperature of 50°C. The Q Exactive HF was operated in a data dependent mode with a survey scan range of 300-1750 m/z and a resolution of 60’000 at m/z 200. MaxQuant software (version 1.5.3.54) was used to analyze MS raw files[57]. MS/MS spectra were searched against the human Proteome Uniprot FASTA database and a common contaminants database (247 entries) by the Andromeda search engine[58].

### Immunofluorescence and confocal microscopy

Between 4 to 5×10^4^ cells were seeded on 12 mm glass cover slips in a 24 well plate and cultured for 1 day before the experiment. Keratinocytes were treated with LecB for the indicated concentrations and time points. Surface or whole cell staining was then performed. For surface staining, NHKs were fixed with 4 % (w/v) formaldehyde (FA) for 10 min and quenched with 50 mM ammonium chloride for 5 min. Cells were blocked with 3 % (w/v) of bovine serum albumin (BSA) in PBS and incubated overnight with the primary antibody diluted in blocking solution (see the details for the used antibodies in table S2). Cells were then washed in PBS and incubated for 30 min with the secondary dye-labelled antibody. For whole cell staining, cells were fixed with 4 % (w/v) FA and quenched, as previously described. When methanol fixation was used instead of FA, cells were incubated with ice cold methanol for 10 min and rinsed twice with PBS. After fixation, 10 min incubation with 0.1 % (v/v) Triton X-100 in PBS was performed. Cells were blocked and subsequently stained with the primary and secondary antibody, as previously described, but using blocking solution supplemented with 0,05 % Triton X-100.

Nuclei were additionally stained with DAPI diluted in PBS with 0.1 % (v/v) Triton X-100. After three additional washes, glass cover slips were mounted with mowiol and imaged using an A1R Nikon confocal microscopy system with a 60x oil immersion objective (NA = 1.49). Z-stacks of at least 3 different areas per condition were acquired and analysed with Fiji ImageJ 1.0 software. Coloc2 Fiji’s plugin was employed for colocalisation analysis.

### Western blot analyses

NHKs were seeded in a 12 well plate at a density of 1.3×10^5^ per well and stimulated with LecB for the indicated time points. After the treatment, cells were washed with PBS and lysed in RIPA buffer (20 mM Tris-HCl, pH 8, 0.1 % (w/v) SDS, 10 % (v/v) glycerol, 13.7 mM NaCl, 2 mM EDTA and 0.5 % (w/v) sodium deoxycholate) in water supplemented with protease and phosphatase inhibitors. Protein concentration was measured using BCA assay kit (Thermo Scientific) and normalised. Samples were then separated via SDS-PAGE and subsequently transferred onto a nitrocellulose membrane. Membranes were blocked in 3 % (w/v) BSA or 5 % (w/v) milk powder in Tris-buffered saline (TBS) supplemented with 0.1 % (v/v) Tween 20 and incubated with the primary and horseradish peroxidase secondary antibodies in blocking solution (see antibody list in Table S2).

Protein bands were visualised via chemiluminescence reaction using the Fusion-FX7 Advance imaging system (Peqlab Biotechnology). A densitometric analysis of at least 3 independent experiments was carried out using FIJI ImageJ 1.0 software.

### Cell viability assays

To assess cytotoxicity and morphological changes upon treatment with LecB, NHKs were seeded in a 24 well plate at a density of 5×10^4^ per well and treated for 24 h with different LecB concentrations ± fucose (30mM). Images were acquired using Evos FL Cell Imaging Systems using 20x objective (NA = 0.45).

MTT cell growth assay kit (Merck) was additionally employed to quantify cell viability upon LecB treatment. To this purpose cells were grown into a 96 well plate and stimulated for 24 h with the indicated LecB concentrations ± L-fucose. The MTT solution was added for 4 h at 37°C with 5 % CO_2_ and the absorbance at 570 nm was measured.

### Ultrastructural analyses

For conventional transmission electron microscopy, NHKs were treated or not with 5 µg/ml of LecB, for the indicated time periods. An equal volume of double strength fixative (4% paraformaldehyde, 4% glutaraldehyde in 0.1 M sodium cacodylate buffer, pH 7.4) was then added to the cells for 20 min at room temperature, prior to fixing the cells with one volume of single strength fixative (2% paraformaldehyde and 2.5% glutaraldehyde in 0.1 M sodium cacodylate buffer, pH 7.4) for 2 h at room temperature. After washes with cacodylate buffer (pH 7.4), cells were then scraped and embedded as previously described [59]. Ultra-thin 70-nm sections were cut using a Leica EM UC7 ultra microtome (Leica Microsystems, Wetzlar, Germany) and stained with uranyl acetate and lead citrate as previously described [59].

For the IEM analyses, NHKs were treated for 12 h with biotinylated LecB and fixed by addition of 4% PFA and 0.4% glutaraldehyde in 0.1 M phosphate buffer (pH 7.4) in an equal volume to the DMEM medium, before to be incubated for 20 min at room temperature. Cells were subsequently fixed in 2% PFA and 0.2% glutaraldehyde in 0.1 M phosphate buffer (pH 7.4) for 3 h at room temperature and then embedded for the Tokuyasu procedure before cutting ultrathin cryo-sectioning and immuno-gold label them, as previously described (Slot and Geuze, 2007). Biotinylated LecB was detected using an anti-biotin rabbit antibody (Rockland, 100-4198). The labelling was revealed with protein A conjugated with 10 nm gold (CMC, Utrecht, NL). As labelling control, NHKs were incubated with not biotinylated LecB. No immuno-gold labelling was detected in this sample. Cell sections were analysed using a CM100bio TEM (FEI, Eindhoven, Netherlands).

### Immunofluorescence of chronic human wounds

Fixed sections of human chronic wounds were kindly provided by A. Nyström (Department of Dermatology, Medical Centre, Albert-Ludwigs-University Freiburg). Antigens were retrieved with 0.05 % pronase (Sigma). Sections were blocked with 3 % (w/v) BSA in PBS and probed overnight with an anti-LecB or anti*-Pseudomonas aeruginosa* antibody. After 2 washing steps with PBS supplemented with 0.1 % Tween 20, the sections were stained with a dye-labelled secondary antibody, counterstained with DAPI and mounted in fluorescence mounting medium (Dako). Images were acquired using Zeiss Axio Imager A1 fluorescence microscopy.

### Statistics

All data were obtained from at least 3 independent experiments and are shown as the means ± standard error mean (SEM).

Statistical analysis was performed using GraphPad Prism software. One-way or two-way ANOVA were chosen to assess significance in experiments with multiple conditions. Otherwise, when experimental data for one condition were compared to the relative control condition, multiple t-test was applied.

A p-value < 0.05 was considered as statistically significant. * denotes p < 0.05; ** denotes p < 0.01; *** denotes p < 0.001; **** denotes p < 0.0001; ns denotes not significant.

## Acknowledgments

The research group of W.R. was supported by the German Research Foundation (EXC 294, GSC-4 and RO 431/2-1), the Ministry of Science, Research and the Arts Baden-Württemberg (Az: 33-7532.20), and by a starting grant from the European Research Council (Programme “Ideas”, ERC-2011-Stg 282105-lec&lip2invade). F.R. is supported by ZonMW VICI (016.130.606), ZonMW TOP (91217002), ALW Open Programme (ALWOP.310) and Marie Skłodowska-Curie Cofund (713660) and Marie Skłodowska Curie ETN (765912) grants. M.M. is supported by an ALW Open Programme (ALWOP.355). We appreciated the support by the BIOSS Toolbox, especially P. Salavei, in the purification of the bacterial lectins.

## Author contributions

A.L. and W. R. designed this interdisciplinary study. The experiments were conducted by A.L. and M.M. under the supervision of W.R. C.G. and S.K. helped in the stainings of tissue samples. Mass spectrometry measurements were performed by T.W. under the supervision of R.G. Furthermore, S.K., I.W., J.C. and R.T. provided preliminary data. D.K. identified patients with *P.* aeruginosa infected chronic wounds and provided the human skin samples. A.L. performed data analysis and wrote the manuscript. M.M., F.R. and A.N. provided experimental advice and expertise. W.R. supervised the study.

## Conflict of interest

The authors declare no conflict of interest.

---

**Table S1.** List of coprecipitated growth factor receptors identified by mass spectrometry analysis.

| **-LogP** | **Difference**<br>**(untreated -**<br>**treated)** | **Protein** | **Gene**<br>**name** |
| --- | --- | --- | --- |
| 4,96 | -8,06 | Cation-independent mannose-6-phosphate<br>receptor | IGF2R |
| 3,70 | -7,95 | Insulin-like growth factor 1 receptor | IGF1R |
| 3,52 | -6,94 | Hepatocyte growth factor receptor | MET |
| 5,08 | -3,81 | Epidermal growth factor receptor | EGFR |

**Table S2.** List of primary antibodies used.

| target | species | supplier | catalog number | dilution (application) |
| --- | --- | --- | --- | --- |
| P-AMPK | rabbit | Cell signaling | 2535 | 1:1000 (WB) |
| AMPK | rabbit | Cell signaling | 2532 | 1:500 (WB) |
| β-actin | mouse | Sigma Aldrich | A5316 | 1:2000 (WB) |
| Cyclin D1 | rabbit | Cell signaling | 2978 | 1:1000 (WB) |
| EEA1 | rabbit | Cell signaling | 3288 | 1:100 (IF) |
| EGF-1R | rabbit | Santa Cruz | Sc-03 | 1:500 (WB) |
| GAPDH | rabbit | Sigma Aldrich | G9545 | 1:2000 (WB) |
| IGF-1R | rabbit | Cell signaling | 9750 | 1:100 (IF)<br>1:1000 (WB) |
| IGF-1R | mouse | Abcam | ab16890 | 1:100 (IF, surface staining) |
| P-IGF-1R (Y1331) | rabbit | Cell signaling | 3021 | 1:1000 (WB) |
| P-IGF-1R (Y1335/1336) | rabbit | Cell signaling | 3024 | 1:1000 (WB) |
| LAMP1 | rabbit | Cell signaling | 9091 | 1:100 (IF) |
| LC3b | rabbit | Abcam | ab48394 | 1:100 (IF)<br>1:1000 (WB) |
| LecB | rabbit | Eurogentec | own production | 1:2000 (WB)<br>1:50 (IF) |
| P-p44/42 MAPK (ERK1/2) | rabbit | Cell signaling | 4370 | 1:1000 (WB) |
| p44/42 MAPK (ERK1/2) | rabbit | Cell signaling | 4965 | 1:1000 (WB) |
| P-mTOR | rabbit | Cell signaling | 2983 | 1:1000 (WB) |
| mTOR | rabbit | Cell signaling | 5536 | 1:1000 (WB) |
| *P. aeruginosa* | rabbit | Abcam | ab68538 | 1:50 (IF) |
| Rab9A | rabbit | Cell signaling | 5118 | 1:100 (IF) |
| Rab11 | rabbit | Cell signaling | 5589 | 1:100 (IF) |
| $\alpha/\beta$-tubulin | mouse | Cell signaling | 2148 | 1:2000 (WB) |
| ubiquitin | mouse | Cell signaling | 3936 | 1:3000 (WB) |
| P-ULK1 (S757) | rabbit | Cell signaling | 14202 | 1:1000 (WB) |
| P-ULK1 (S555) | rabbit | Cell signaling | 5869 | 1:1000 (WB) |

**Table S3.** List of secondary antibodies used.

| target | tag | supplier | catalog number | dilution<br>(application) |
| --- | --- | --- | --- | --- |
| anti-mouse | HRP | Cell signaling | 7076 | 1:1000/1:2000<br>(WB) |
| anti-rabbit | HRP | Cell signaling | 7074 | 1:1000/1:2000<br>(WB) |
| anti-mouse | Cy3 | Jackson<br>Immunoresearch | 715-166-150 | 1:200 (IF) |
| anti-rabbit | Cy3 | Jackson<br>Immunoresearch | 711-166-152 | 1:200 (IF) |
| anti-mouse | DyLight 650 | Thermo Fisher | 84545 | 1:200 (IF) |
| anti-rabbit | DyLight 650 | Thermo Fisher | 84546 | 1:200 (IF) |
| anti-rabbit | Alexa Fluor 647 | Thermo Fisher | A-21245 | 1:200 (IF) |

**SI Appendix Fig. S1.**
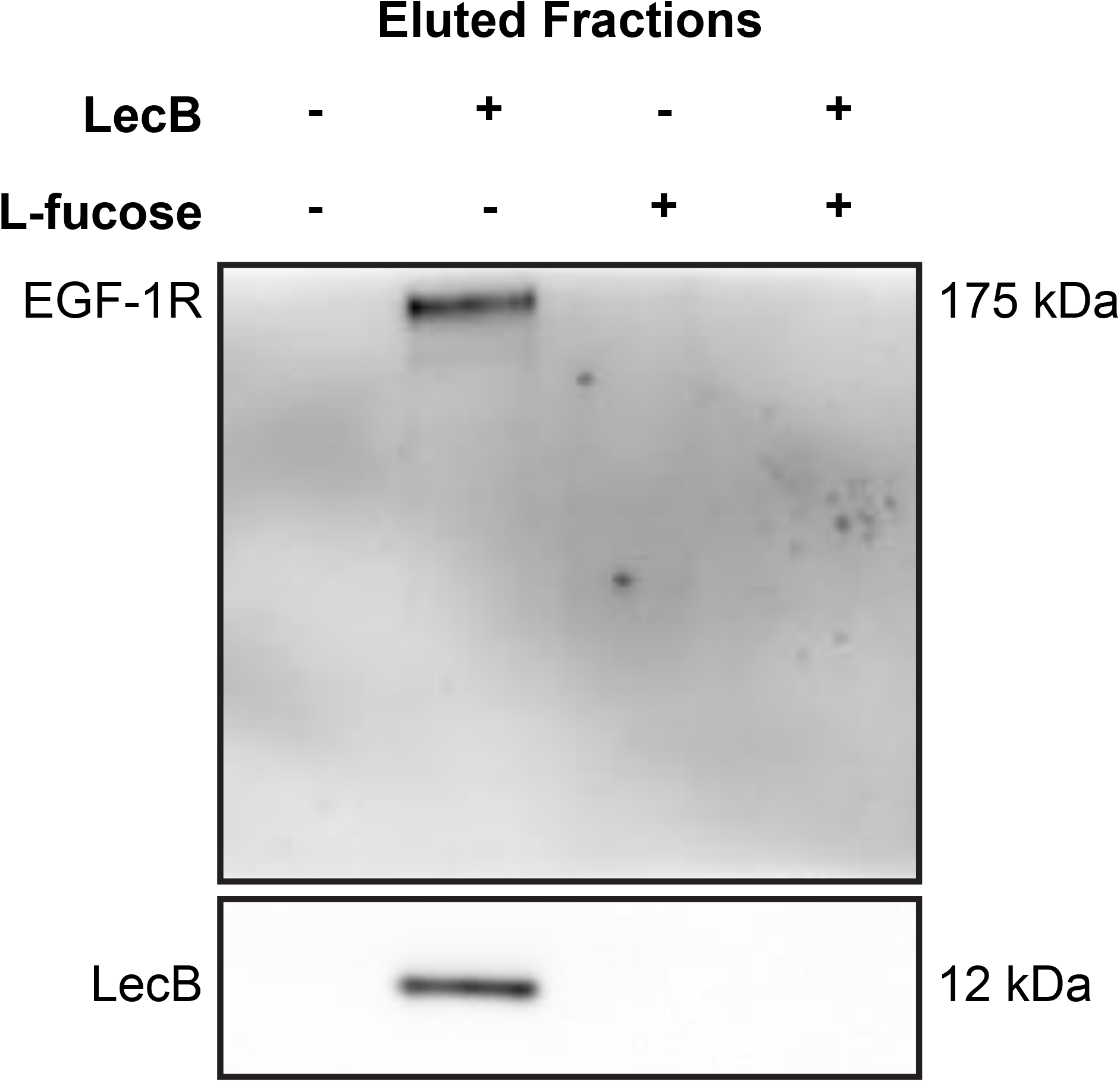
Western Blot of eluted samples from pull-down assay using biotinylated LecB (5 μg/mL) with or without 30 mM L-fucose. Membranes were probed for LecB and EGF-1R.

**SI Appendix Fig. S2.**
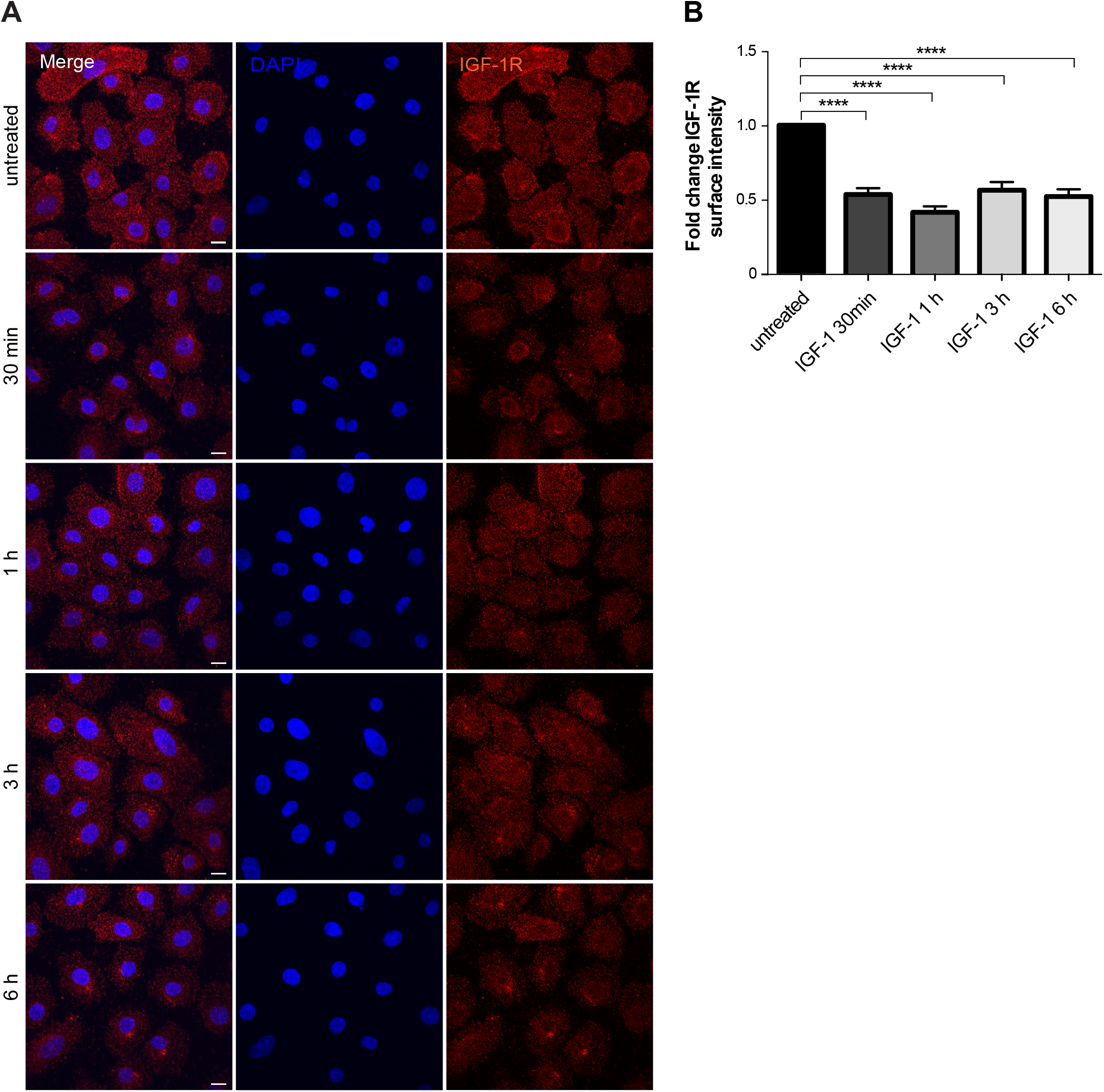
(A) Surface staining of IGF-1R (red) upon stimulation of keratinocytes with 100 ng/mL IGF-1 for the indicated times. Maximum intensity projections are depicted. Nuclei are indicated by DAPI (blue). (B) Quantification of IGF-1R surface intensity. Bars show the mean value ± standard error mean of n = 3 independent experiments. **** denotes p < 0.0001.

**SI Appendix Fig. S3.**
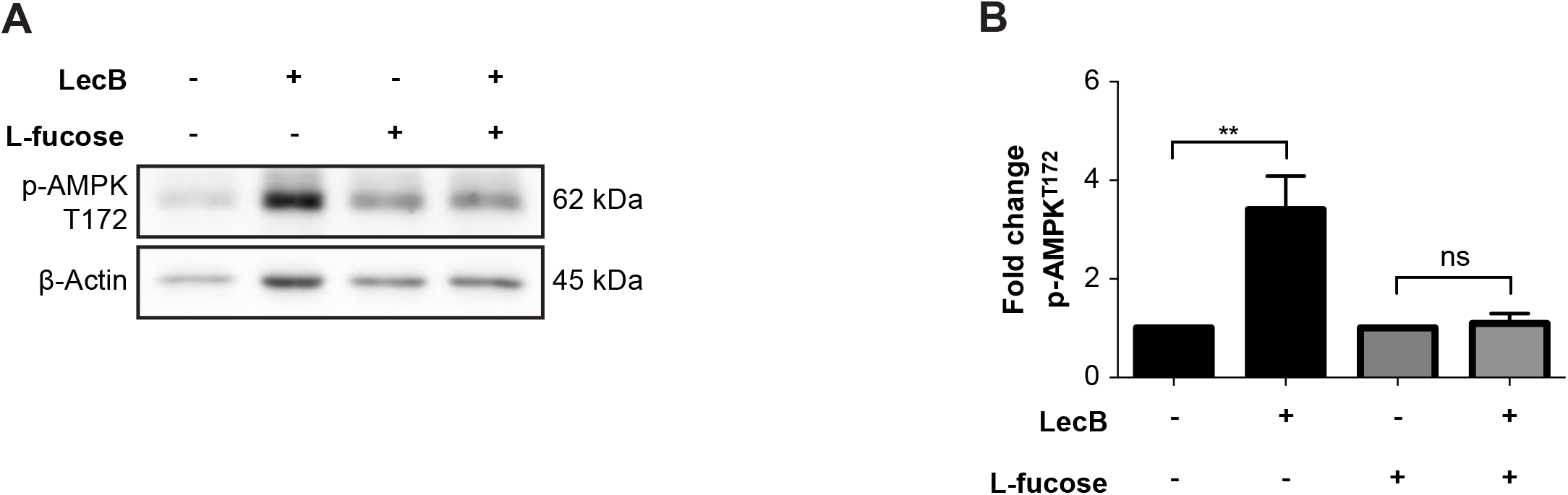
(A, B) Whole cell lysates were analysed by Western Blot to detect levels of pAMPK upon 4 h incubation with LecB (5 μg/mL) ± 30 mM L-fucose. The phosphorylated protein levels were normalised to actin. The graph reports the mean value ± standard error mean of N = 3 independent experiments. ** denotes p < 0.01; ns denotes not significant.

**SI Appendix Fig. S4.**
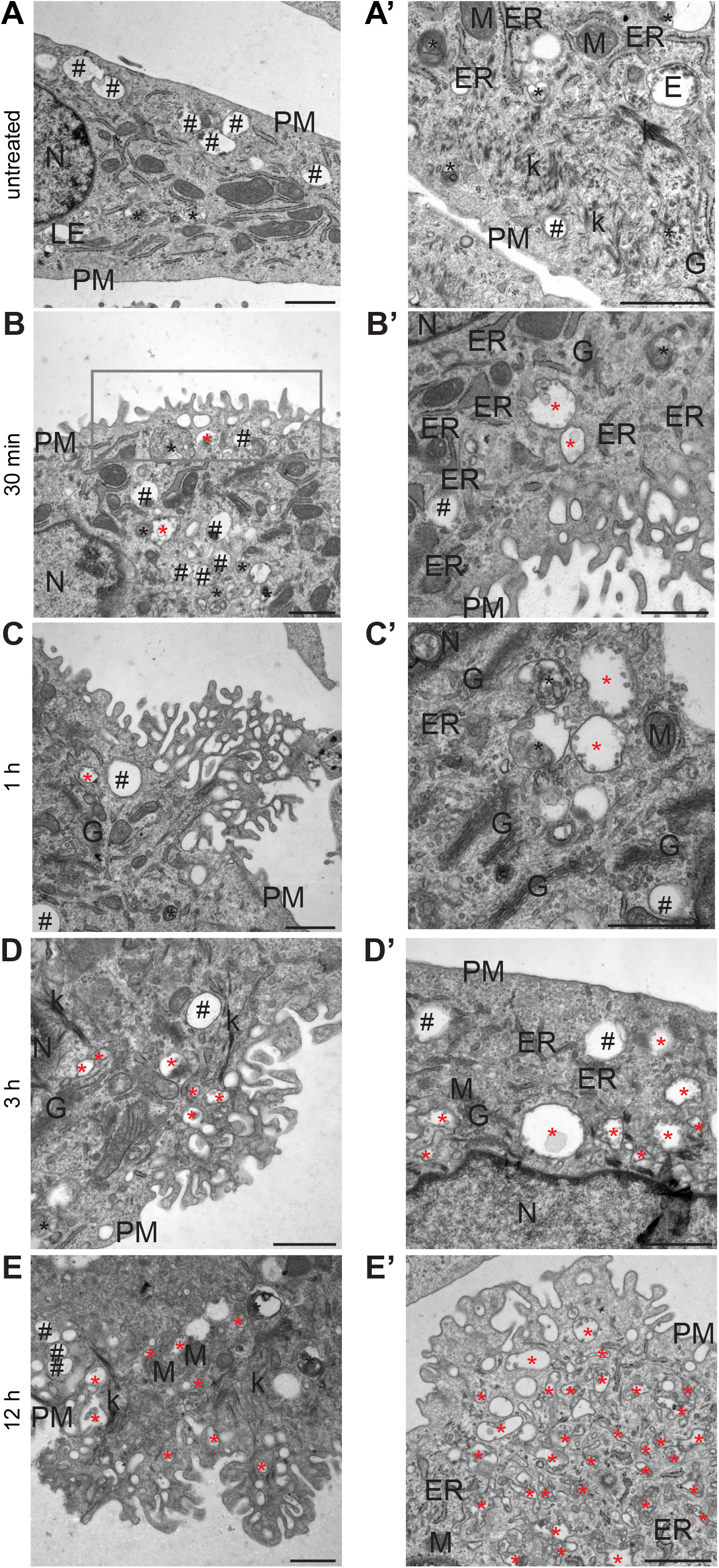
LecB containing vesicles originate from ruffle-like regions at the plasma membrane. (A-E) Representative electrographs of keratinocytes treated with 5 μg/mL LecB for the indicated time points and processed for conventional EM. (A’-E’) Additional area for each indicated condition with higher magnification. Black hashtags point at category 1 vacuoles, while black and red asterisks indicate category 2 and category 3 vacuoles, respectively. Rectangular box highlights ruffle-like regions. Scale bars: 1 μm E: endosome; ER: endoplasmic reticulum; k: keratin; G: Golgi apparatus; M: mitochondrion; N: nucleus; PM: plasma membrane.

**SI Appendix Fig. S5.**
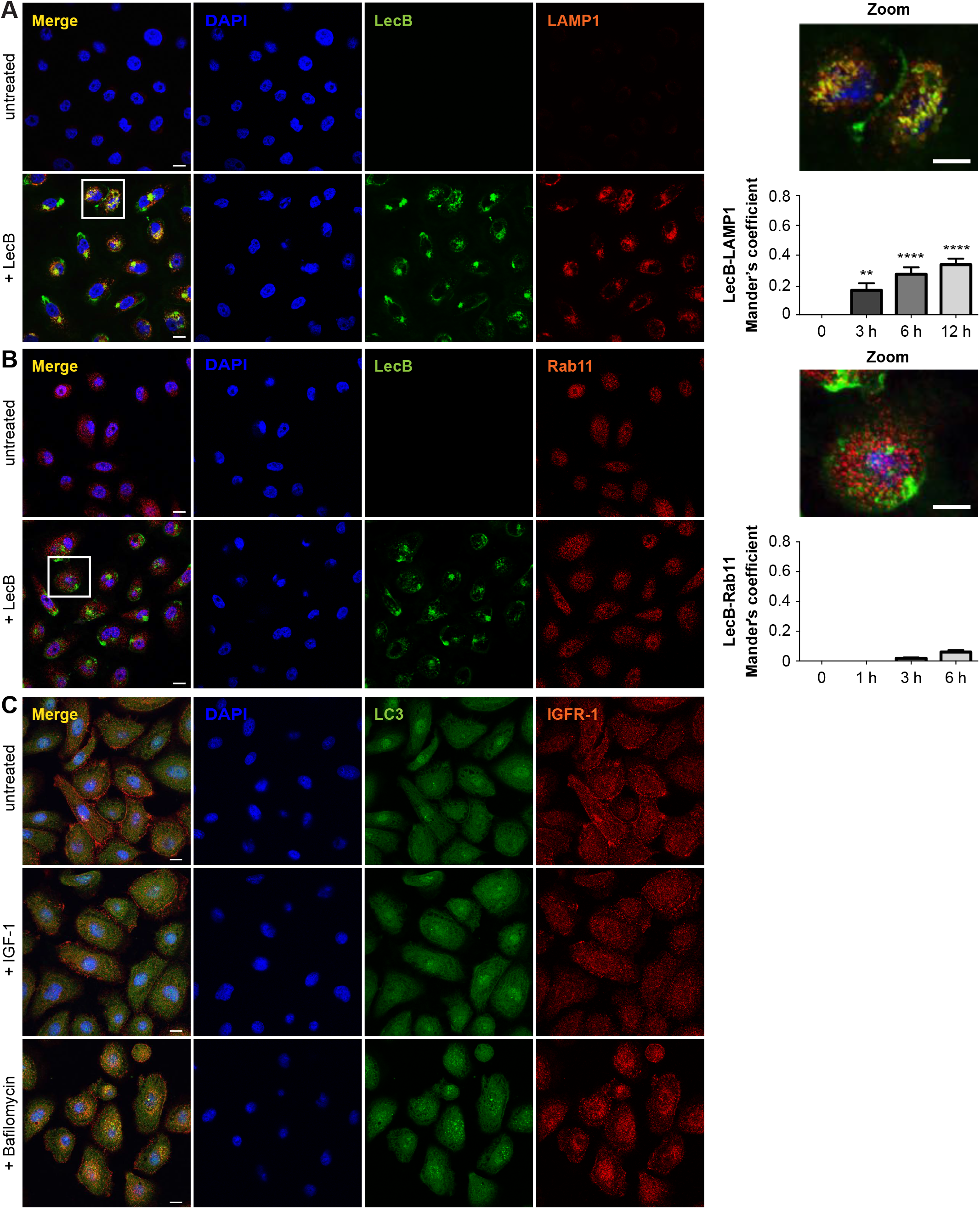
(A, B) Confocal micrographs of NHKs stimulated with 5 μg/mL of fluorescently labelled LecB (green) permeabilised and stained for (A) LAMP1 and (B) Rab11 (both in red). Panels indicated as “+ LecB” refer to 12 h incubation. Note that LAMP1 signal in the untreated panel is not visible due to the high difference in intensity between untreated and treated conditions. Scale bar: 10 μm. Graphs right to the panels show the quantification of Mander’s overlap coefficients between LecB and LAMP1 and LecB and Rab11. Error bars indicate means ± standard error mean of N = 3 independent experiments. ** denotes p < 0.01; **** denotes p < 0.0001. (C) Representative confocal images depicting NHKs treated with 100 ng/mL IGF-1 or 100 nM bafilomycin for 12 h and stained for LC3 (pink) and IGF-1R (red). N = 3.

**SI Appendix Fig. S6.**
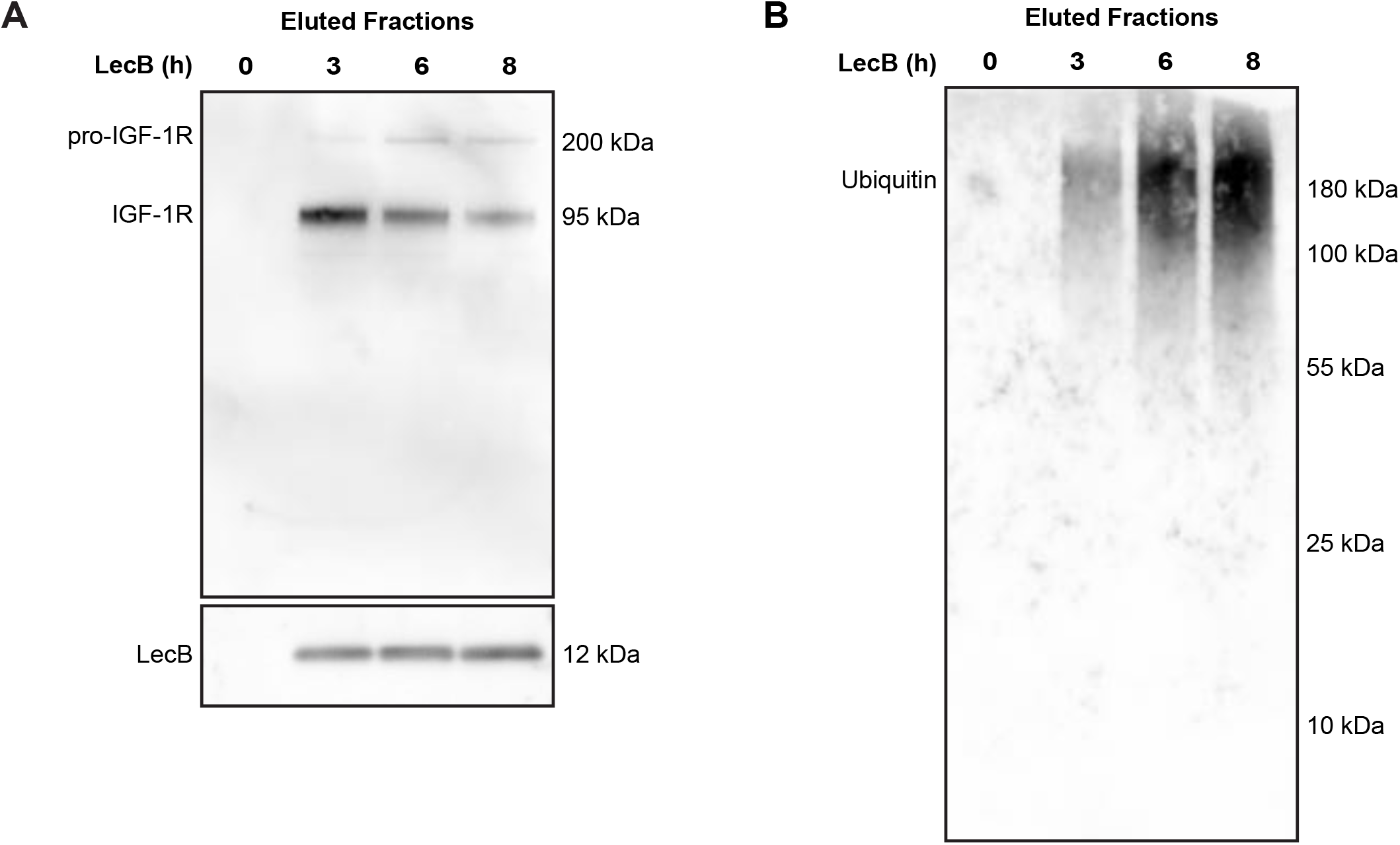
(A, B) Western Blot of eluted samples from time series pull-down assay using biotinylated LecB (5 μg/mL). Membranes were probed for LecB, IGF-1R (A) and ubiquitin (B).

**SI Appendix Fig. S7.**
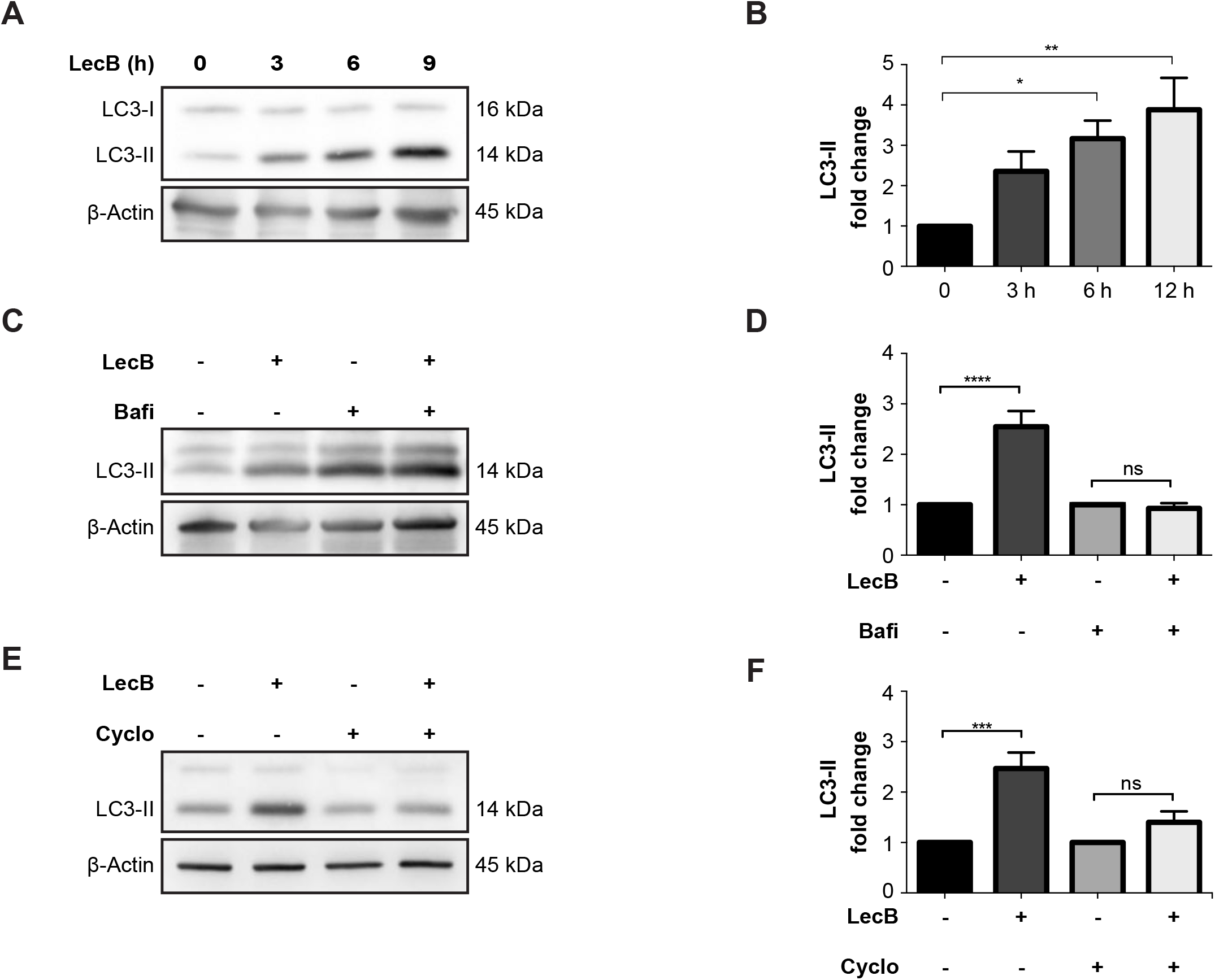
(A, B) NHKs were incubated with LecB (5 μg/mL) for the indicated time points and whole cell lysates were immunoblotted for LC3 and β-actin (A). The levels of LC3-II were normalised to β-actin and depicted as fold change increase to the untreated condition (B). Error bars indicate means ± standard error mean of N = 5 independent experiments. (C-F) Representative blots and relative quantifications. Cells were treated with 5 μg/mL of LecB for 12 h ± 200 nM bafilomycin (Bafi) or ± 20 μg/mL cycloheximide (Cyclo). Whole cell lysates were probed for LC3 and β-actin was used as loading control. Graphs indicate the fold change of LC3-II levels, expressed as means ± standard error mean of N = 6 (C, D) and N = 4 (E, F) independent experiments. *denotes p < 0.05; ** denotes p < 0.01; *** denotes p < 0.001; **** denotes p < 0.0001; ns denotes not significant.

